# Parapatric speciation with recurrent gene flow of two sexual dichromatic pheasants

**DOI:** 10.1101/2021.11.20.469217

**Authors:** Zheng Li, Jie Zhou, Minzhi Gao, Wei Liang, Lu Dong

## Abstract

**Background:** Understanding speciation has long been a fundamental goal of evolutionary biology. It is widely accepted that speciation requires an interruption of gene flow to generate strong reproductive isolation between species, in which sexual selection may play an important role by generating and maintaining sexual dimorphism. The mechanism of how sexual selection operated in speciation with gene flow remains an open question and the subject of many research. Two species in genus *Chrysolophus*, Golden pheasant (*C. pictus*) and Lady Amherst’s pheasant (*C. amherstiae*), which both exhibit significant plumage dichromatism, are currently parapatric in the southwest China with several hybrid recordings in field.

**Methods:** In this research, we estimated the pattern of gene flow during the speciation of two pheasants using the Approximate Bayesian Computation (ABC) method based on the multiple genes data. With a new assembled *de novo* genome of Lady Amherst’s pheasant and resequencing of widely distributed individuals, we reconstructed the demographic history of the two pheasants by pairwise sequentially Markovian coalescent (PSMC).

**Results:** The results provide clear evidence that the gene flow between the two pheasants were consistent with the prediction of isolation with migration model for allopatric populations, indicating that there was long-term gene flow after the initially divergence (ca. 2.2 million years ago), and further support the secondary contact when included the parapatric populations since around 30 ka ongoing gene flow to now, which might be induced by the population expansion of the Golden pheasant in late Pleistocene.

**Conclusions:** The results of the study support the scenario of speciation between Golden pheasant (*C. pictus*) and Lady Amherst’s pheasant (*C. amherstiae*) with cycles of mixing-isolation-mixing due to the dynamics of natural selection and sexual selection in late Pleistocene that provide a good research system as evolutionary model to test reinforcement selection in speciation.

## Background

The fascination that evolutionary biologists have long had with speciation has increased in recent decades because of a growing understanding of the complex mechanisms that shape biodiversity. Allopatric speciation driven by geographical isolation, implying in the absence of gene flow, has been the standard model to interpret the classical speciation progress (Mayr 1942; Coyne & Orr 2004). However, allopatric speciation models without gene flow seem oversimplified since the recurrence of land bridges during glacial periods of Pleistocene has provided opportunities for gene exchange between populations (Rohling et al. 1998, Lambeck and Chappell 2001). Since more empirical examples of divergence in sympatry or parapatry were reported, it is widely accepted that speciation can occur in the presence of gene flow (Rice and Salt 1988; Bolnick and Fitzpatrick 2007; Nosil 2008). To understand how speciation may proceed in the face of gene flow, one must understand the pattern and role of gene flow in the speciation process.

Many researches so far has focus on the speciation process in monomorphic species of birds, such as Hwameis (Li et al. 2010), Crows (Poelstra et al. 2014), Darwin’s Finches (Lamichhaney et al. 2015) and Bamboo Partridges (Wang et al. 2021), most of that revealed the important impacts of disruptive selection or assortative mating that promote the speciation with gene flow. However, the research on evolutionary mechanisms and speciation in sexual dichromatic species has been lagging behind (Servedio 2011). Compared with sexual monomorphic species, sexual dichromatic species might be exposed to a greater selection, namely Fisherian sexual selection, which plays an important role in driving population divergence as an essential driver of pre-zygotic isolation. However, there are only some theoretical researches based on model simulation (Servedio 2012; Servedio and Bürger 2014; Servedio 2016) for the pattern and role of gene flow in speciation under Fisherian sexual selection (but see Dong et al. 2013 for a case study in birds). More empirical researches are still needed to support theoretical expectations.

To examine the pattern and role of gene flow in speciation between sexual dichromatic species, we focus specifically on two species in genus *Chrysolophus*, golden pheasant (*C. pictus*) and Lady Amherst’s pheasant (*C. amherstiae*) which both exhibit significant plumage dichromatism. The males of golden pheasant and Lady Amherst’s pheasant have extreme phenotypic differences in their plumage color, but the females are very similar, both with a duller mottled brown plumage. The golden pheasants are mainly distributed in the mountains of central and west China from the Qinling Mountains in the north to the Minshan Mountain in the west, while the Lady Amherst’s pheasants are mainly distributed in the Yunnan-Guizhou Plateau and the western and southern edge of the Sichuan Basin (Fig. 1). These two pheasants are currently parapatric in southwest China with several hybrid recordings in west Sichuan (He and Wei 1993; Shi et al. 2018) and northeast Yunnan (Deng 1974), and even a hybrid recording in a painting attributed to Emperor Huizong of the Song Dynasty around one thousand years ago (Peng et al. 2016). These ongoing hybridization records implied incomplete pre-zygotic reproductive isolation in the sympatric zone. And the captive observation also revealed incomplete post-zygotic reproductive isolation in the F1 hybrids and backcross hybrids with higher proportion of the male hybrids which corresponded to the prediction of Haldane’s rule (Arrieta et al. 2013).

**Fig. 1.**
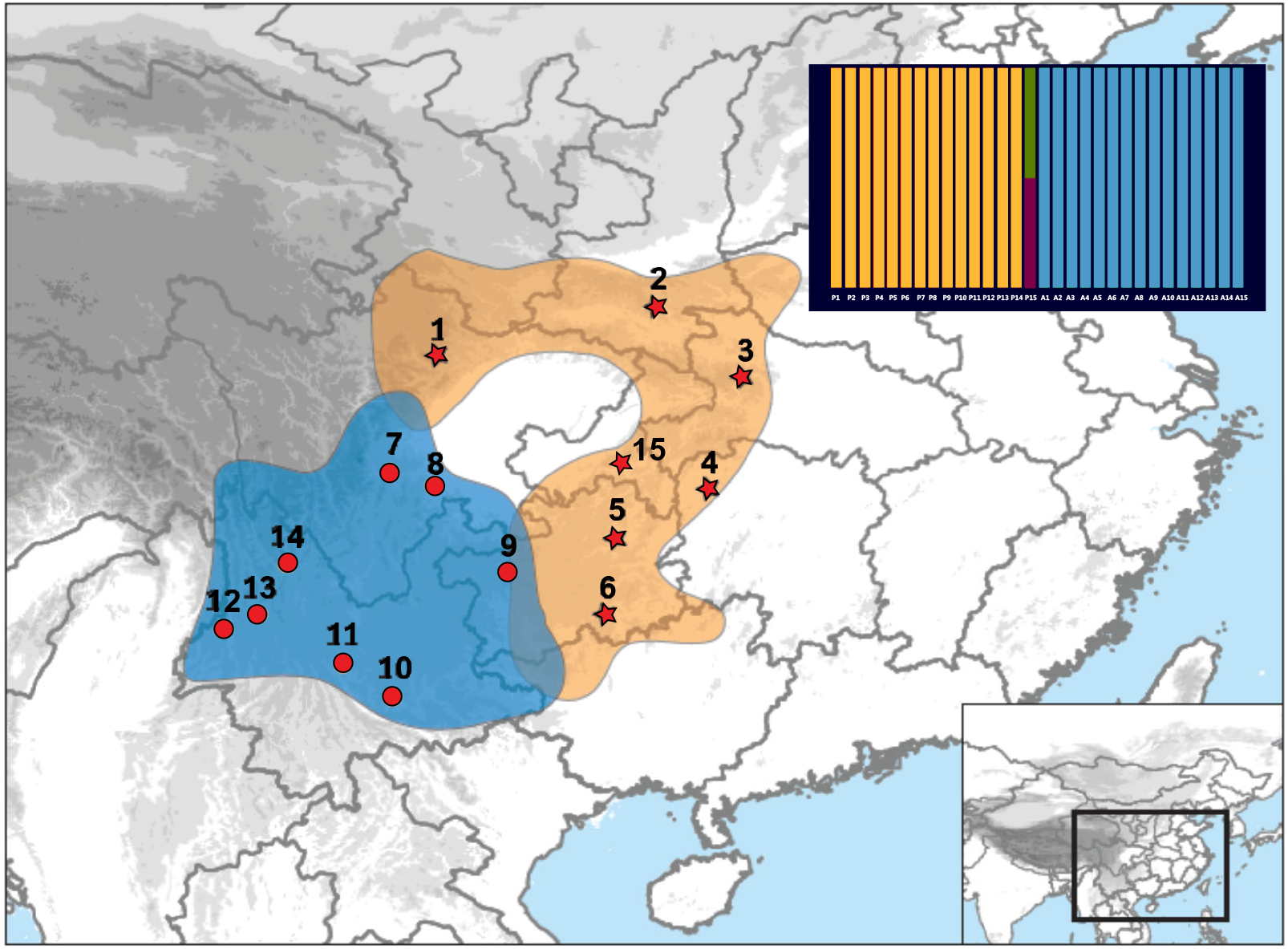
The distribution and sampling locations of *C. pictus* (orange zone and red star) and *C. amherstiae* (blue zone and red circle)

The niche model analysis explained the divergence history since the last interglacial (LIG) of the two pheasant species. The significantly different ecological niches occupied by the *C. pictus* and *C. amherstiae* according to current distribution and the isolated glacial shelters in the late Pleistocene may promote the allopatric speciation of the two species (Lyu et al. 2015). However, without genetically based coalescence analysis, this model is limited to reconstructing the divergent history before the segregation of the two species before the LIG and distinguishing the mode of the speciation process, thus what happened since the divergence of *C. pictus* and *C. amherstiae* cannot be estimated. The ecological niches between the two *Chrysolophus* species were different in the late Pleistocene, allopatric with distribution gap, and re-parapatric in/after LGM that may be induced by expansion of the golden pheasant, that provide a good research system for studying the speciation with gene flow under both sexual selection and natural selection.

In this study, we combined the multi-locus data from the two closely related species of *Chrysolophus* and a *de novo* genome of *C. amherstiae* with several resequencing individuals from two pheasant species to achieve two main goals. First, we aimed to detect the candidate hybrids with phylogenetic analysis and further explore the mode of gene flow in speciation of *C. pictus* and *C. amherstiae* through Approximate Bayesian Computation (ABC) analysis (Beaumont et al. 2002). Second, we employed pairwise sequentially Markovian coalescent (PSMC) analyses (Li and Durbin 2011) to inspect the evolutionary history between *C. pictus* and *C. amherstiae*, basing on the draft genome and re-sequencing data. These analyses allowed us to explore the pattern and role of gene flow in speciation of sexual dichromatic species and provide a better understanding of how speciation with gene flow proceed.

## Methods

### Sampling, amplification and sequencing

A total of 38 individuals (23 individuals of *C. pictus*, 15 individuals of *C. amherstiae*) from 15 locations were sampled across the geographic range of *C. pictus* and *C. amherstiae*, 30 of them were used for ABC analysis and 19 for PSMC analysis (Fig. 1; Table S1). One Temminck’s Tragopan *Tragopan temminckii*, collected from Shimian County, Sichuan Province, was used as the outgroup in phylogenetic analysis. DNA was extracted from ethanol-preserved muscle, blood or feather using TIANamp Genomic DNA kit (Tiangen Biotech Ltd., Beijing) following the manufacturer’s protocol and dissolved in 100 μl of TE buffer. In total of 37 exon loci (Table S2) including 34 unlinked autosomal loci and two Z-linked loci developed by Liu et al. (2018) and the mitochondrial cytochrome b (*cyt* b) gene were amplified by polymerase chain reaction (PCR). The Touchdown PCR (Don et al. 1991) was performed in a PTC-220 PCR thermal cycler system (Bio-Rad, USA). The 40 μl reaction for PCR contained 2.4 μl template DNA, with 20ul mixed concentrations of 2 × ExTaq buffer (Takara, Japan), 0.8ul of each forward and reverse primer (10uM), and 16ul ddH2O. The initial denaturation at 94°C for 3 min, followed by 20 cycles of 94°C for 30 s, 62–52 °C (decreasing the annealing temperature by 0.5 °C per cycle) for 30 s and 72 °C for 1min, and another 15 cycles of 94°C for 30 s, 52 °C for 30s and 72 °C for 1min, ended with a final extension at 72°C for 10 min. PCR products were sequenced bidirectionally using the 3730XL series platform Sequencer (Applied Biosystems, USA).

### Sequences analysis

The obtained sequences were edited using the software CodonCode Aligner (CodonCode Corporation, Dedham, MA, USA), aligned with Clustal W (Thompson et al. 1994) in MEGA v5.10 (Tamura et al. 2013) and removed the sequences with multiple deletions. Haplotypes of the diploid nuclear sequences were separated using DNASP v5.10.01 (Librado and Rozas 2009) based on Markov chain Monte Carlo (MCMC) simulation approach with 500 bootstrap replicates and the results of the first 100 times were discarded. Due to the small sample size, haplotypes of the diploid nuclear sequences of *T. temminckii* were manually separated. The phased nuclear sequences and cytb sequences were aligned between *C. pictus, C. amherstiae* and *T. temminckii* with Clustal W (Thompson et al. 1994) in MEGA v5.10 (Tamura et al. 2013) and saved as fasta format for subsequent analyses.

We tested the neutrality of each gene using Hudson-Kreitman-Aguade (HKA) test (Hudson et al. 1987) and Tajima’s D (Tajima 1989). We calculated intra- and interspecific summary statistics for DNA polymorphisms of each gene (Table S3) by DnaSP v5.10.01 (Librado and Rozas 2009) for the subsequent ABC analysis (Beaumont et al. 2002). No significant λ^2^(HKA) and Tajima’s D were observed in all genes except for the Tajima’s D of the mitochondrial cytb gene (Table S4). Nuclear substitution rate was calculated by taking the average ratio of within group and outgroup pairs and then multiplied it by the rate of 1.035 × 10^−8^ per site per million years (molecular clock for the passerine mitochondrial cytb gene, Weir and Schluter 2008). The mitochondrial cytb gene was not included in the ABC and PSMC analyses because its evolutionary rate highly differs from that of nuclear genes.

### Detection of hybrids

We used the NewHybrids (Anderson and Thompson 2002) to calculate the posterior probability of sampled individuals belonging to parental or hybrid categories based on the genotypic information of the two pheasants. In this study, individuals collected 50 km away without niche similarity from the sympatric zone (Lyu et al. 2015) were used as the parent group to estimate the allele frequencies of individuals collected in the suture zone or apart within 50 km. The dataset was analyzed three times using the uniform prior for both θ and the mixing proportion π. The burn-in periods and numbers of sweeps were set as recommended value in the manual. The mean value was used to infer hybrids between the two pheasants.

To infer the phylogenetic relationship between pheasants sampled, we used the ancestral reconstruction algorithm implemented in BEAST 1.8 (Drummond et al. 2012) based on autosome nuclear genes, Z chromosome genes and cyt b gene respectively. The substitution model of cyt b gene were determined with jModelTest2.1.3 (Darriba et al. 2012) based on the Bayesian information criterion (BIC). Strict clock was set for each gene, the rate of cyt b gene were set to 1.0 and the substitution rate of nuclear genes were estimated relative to the rate of cyt b gene. Length of MCMC chain for the BEAST analysis were set at 1×10^7^ steps and the first 10% steps of each run were discarded as burn-in. Convergence were checked with Tracer 1.5 (Rambaut and Drummond 2009) to make sure the ESS values of main parameters above 200.

### Approximate Bayesian Computation (ABC)

To infer the speciation history of the two *Chrysolophus* species, we used ABC analysis, which has been widely applied to phylogeographic studies (Beaumont et al. 2002; Beaumont 2010; Bertorelle et al. 2010; Duchen et al. 2013; Cornejo-Romero et al. 2017) to statistically test four alternative models of demographic history: (1) Isolation model, no gene flow occurs throughout the whole divergence process. (2) Isolation with migration model (IM model), gene flow continuously occurs after an initial population split. (3) Early gene flow model, gene flow only occurs at the initial stages of divergence. (4) Secondary contact model, gene flow only exists in the late period of divergence. The parameters to be estimated for the four models are as follows: divergence time (T_div_), Effective population size (N_e_) of *C. pictus* (N_cp_) and *C. amherstiae* (N_ca_), N_e_ of the most common ancestor (N_A_). For the model with gene flow, the number of migratory in each generation was estimated: M_1_ represents the number of migrants of *C. pictus*, M_2_ represents the number of migrants from *C. amherstiae*, m_cp_ represents the migration rate of *C. pictus*, and m_ca_ represents the migration rate of *C. amherstiae*.

For each speciation model, we used the software msABC (Pavlidis et al. 2010) to perform multi-locus coalescent simulations for 1×10^6^ times and each divergence time was normalized with 4N_e_. Demographic parameters were randomly sampled from a uniform distribution within the prior ranges. The ranges of the priors for each demographic parameter in all models and the command line for each model used in msABC are listed in Additional file 1: Tables S5 and S6, respectively.

We performed the model selection, and parameter estimation under the most probable model using “abc” package (Csilléry et al. 2012) in R statistical programming environment (R Core Team 2020). With the 1×10^6^ simulations, posterior probabilities for each demographic model were computed using a multinomial logit regression (tolerance level=0.001) (Beaumont et al. 2008). We selected the model with the highest posterior probability as the most probable model and performed 3 × 10^6^ coalescent simulations under the most probable model to calculate the parameters. The logit transformations of parameters were applied to estimate the posterior distributions of parameters under the most probable model by neural network regression (Blum and François 2010). Each simulated dataset was weighted by its Euclidian distance through an Epanechnikov kernel (Jean-Marie et al. 2008), and shown in mean and median (Table 1).

**Table 1.** Population parameters of two datasets under isolation with migration model and secondary contact model estimated by ABC analysis. T_div_: population divergence time; T_2_: termination time of gene flow after secondary contact; M_1_: the number of migrants from *C. amherstiae* to *C. pictus* in each generation. M_2_: the number of migrants from *C. pictus* to *C. amherstiae* in each generation.

|  | Allopatric samples |  |  | All samples |  |  |  |  |  |
| --- | --- | --- | --- | --- | --- | --- | --- | --- | --- |
|  | Isolation with migration model |  |  | Isolation with migration model |  |  | Secondary contact model |  |  |
| | $T_{div}$ | $M_1$ | $M_2$ | $T_{div}$ | $M_1$ | $M_2$ | $T_2$ | $M_1$ | $M_2$ |
| 2.5% HPD | $4.26 \times 10^5$ | 0.184 | 0.254 | $4.57 \times 10^5$ | 0.116 | 0.473 | $0.031 \times 10^5$ | 0.668 | 2.14 |
| Median | $22.8 \times 10^5$ | 0.929 | 1.70 | $19.3 \times 10^5$ | 0.602 | 1.774 | $0.130 \times 10^5$ | 2.92 | 8.97 |
| Mean | $22.3 \times 10^5$ | 1.054 | 2.00 | $18.6 \times 10^5$ | 0.713 | 1.054 | $0.258 \times 10^5$ | 3.22 | 9.44 |
| 97.5% HPD | $38.3 \times 10^5$ | 2.590 | 5.34 | $28.8 \times 10^5$ | 1.880 | 1.316 | $1.206 \times 10^5$ | 7.04 | 19.33 |

Considering that the sympatric populations and hybrids may affect the result of model selection, analyses were conducted on two datasets respectively: (1) Allopatric populations of *Chrysolophus* (excluding the individuals collected from the hybrid zone and 50 km apart from the hybrid zone: site 1, 7 and 9 in Fig. 1); (2) All sampled individuals of two *Chrysolophus* species.

### Whole-genome assembly and mapping of reads

*De novo* sequencing of an individual of one male *C. amherstiae* (collected in Dali, Yunnan, China) was carried out on an Illumina HiSeq 2000 platform, by Novogene Company (Beijing, China). In total of 116.4Gb data was sequenced and assembled by ALLPATHS-LG v41687 (Butler et al. 2008) using default parameters (Table S7).

We resequenced 19 individuals from two *Chrysolophus* species (13 golden pheasants and six Lady Amherst’s pheasants, Table S4) with an average of 35× depth using Illumina NovaSeq 6000 deal-index sequencing libraries by Novogene Company (Beijing, China). The trimmed clean reads of each individual were mapped back onto the assembled reference genome of *C. amherstiae* by BWA-MEM v0.7.12 (Li 2013) using default parameters. We converted SAM files into binary BAM files, sorted the BAM files and removed the PCR repeats using SAMtools v0.1.19 (Li et al. 2009). The final files were used as the input files of PSMC analyses.

### PSMC analyses

By utilizing information from whole genome of a single diploid individual, the PSMC analysis (Li and Durbin 2011) can represent an appropriate method for estimating population size changes deep into the early Pleistocene. We applied the PSMC approach on the genomes of *C. pictus* (13 individuals) and *C. amherstiae* (6 individuals) to reconstruct demographic history of each species. The consensus sequence (in BAM format) for each individual of each species was divided into non-overlapping 100-bp bins, where each bin is scored as heterozygous (‘1’) if there is a heterozygote in the bin, or as homozygous (‘0’) otherwise implemented with the fq2psmcfa command from R package “psmc” and the quality cut off (-q) was set to 20. The psmcfa sequences were taken as the input of the PSMC estimation to deduce the past N_e_ over time. The number of iterations (-N) was set to 25, Tmax (-t) was set to 15, initial mutation/recombination ratio (-r) was set to 5 and the atomic time interval pattern (-p) was ‘4+25*2+4+6’. The psmc file was visualized with the mutation rate (μ) being set to 3.8 × 10^−9^ per site per generation (generation time = 2 years) based on the mean nuclear substitution rate of avian (Zhang et al. 2014).

## Results

### Identification of hybrids

The Bayesian tree demonstrated substantial support for mutual monophyletic clades of the two pheasant species with the exception of incongruent phylogenetic position of one male Golden Pheasant (i.e. SCPW-02) collected near the suture zone (Pingwu, Sichuan) using different kind of molecular markers. The alleles of the male were clustered with the *C. amherstiae* when the autosome nuclear loci were employed to reconstruct the Bayesian phylogenetic relationship (Fig. S1), while were clustered with the *C. pictus* clade reconstructed by the mitochondrial cyt b gene (Fig. S2). When the Z-link alleles were applied, the male alleles were found to be clustered with the two clades, respectively (Fig. S3).

The phylogenetic incongruence was highly supported by the inference from the NewHybrids analyses that the male was detected as the hybrid of F2 (posterior probability = 0.493) or backcross (posterior probability = 0.493) (Fig. 1).

### Evolutionary history with ABC estimation

Since we performed ABC analyses on the two datasets separately, the results of estimation will be presented respectively in the following paragraphs.

For allopatric population of *Chrysolophus*, the best-supported model corresponds to an IM model (posterior probability = 0.97, Fig. 2b), indicating that ongoing gene flow have been existing between *C. pictus* and *C. amherstiae* throughout the whole divergence process. The divergence time of *C. pictus* and *C. amherstiae* was estimated as 2.23 million years ago (Mya), and the gene flow from *C. pictus* to *C. amherstiae* (M_2_ = 2.00) was higher than that from *C. amherstiae* to *C. pictus* (M_1_ = 1.054) during the *Chrysolophus* divergence. The N_e_ of *C. amherstiae* (N_ca_ = 1.191 × 10^5^) was much larger that of *C. pictus* (N_cp_ = 0.617 × 10^5^) and both of them increased compared with the ancestor population size (0.35 × 10^5^, Table S5). All of estimations mentioned above were conducted under the IM model.

**Fig. 2.**
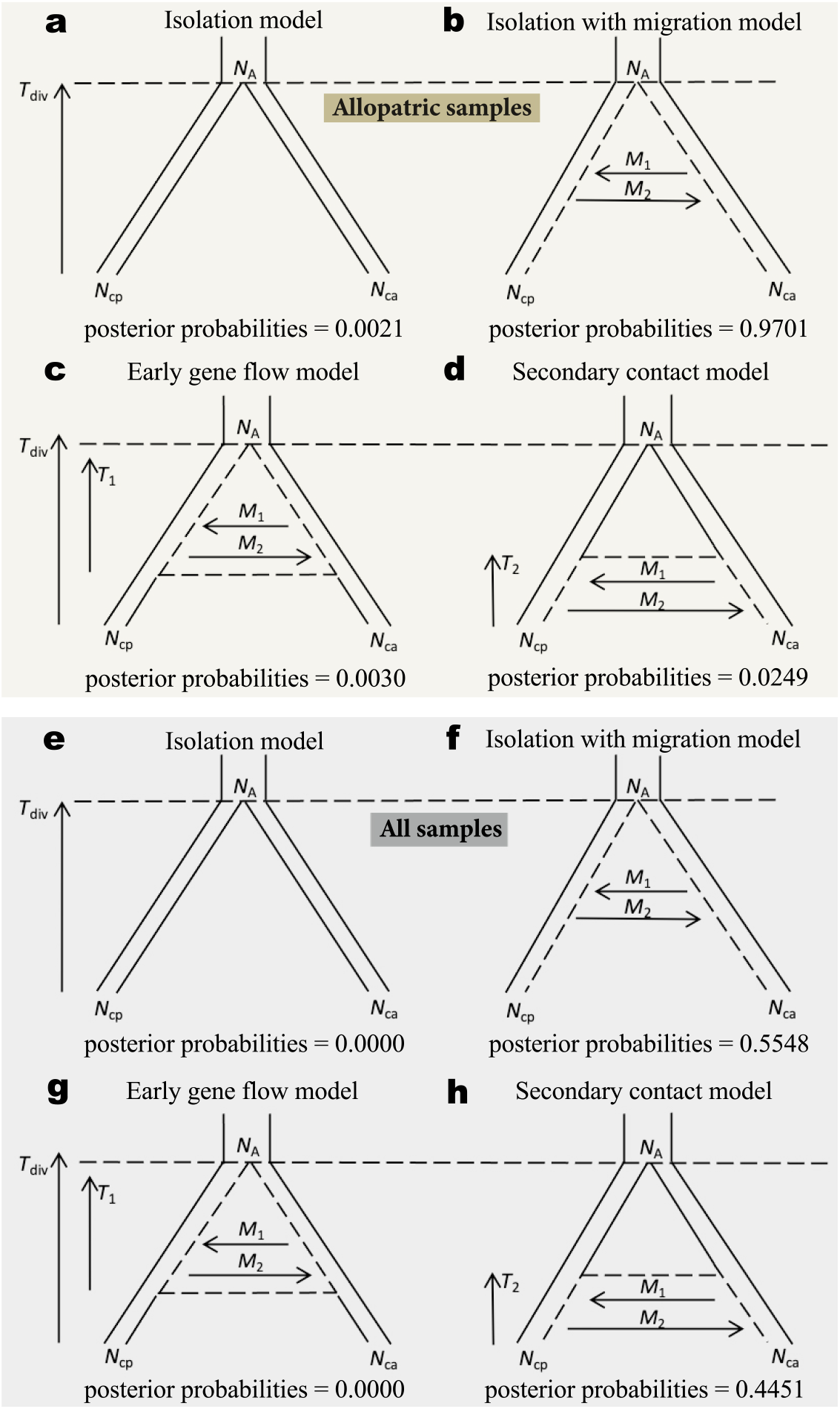
Demographic history scenarios tested in this study, which included posterior probabilities for each model and parameters for divergence time (T_div_), The time when there is gene flow in the early period (T_1_), the time when there is gene flow in the late period (T_2_), ancestral population size (N_A_), *C. pictus* and *C. amherstiae* sizes (N_cp_ and N_ca_, respectively) and gene flow (M_1_ and M_2_) when applicable. For (a to d), samples only include allopatric population of *Chrysolophus*. For (e to h), all sampled individuals of *Chrysolophus*, including individuals collected from hybrid zone.

For all sampled individuals of *Chrysolophus*, the best-supported model is the same IM model (posterior probability = 0.55, Fig. 2f), however the posterior probability of the secondary contact model significantly increased compared with allopatric population estimation (posterior probability = 0.45, Fig. 2h). The divergence time of *C. pictus* and *C. amherstiae* under the best-supported IM model was estimated as 1.86 Mya, and the gene flow of the two pheasants was still asymmetric that gene flow from from *C. pictus* to *C. amherstiae* remains higher (Table 1). We further estimated the parameters of population dynamics under secondary contact model since the posterior probabilities of secondary contact model and isolation with migration model were very close. The initial secondary contact time of the two pheasants is estimated to be about 25,800 years ago, indicating the new hybrid zone was formed after the last glacial maximum. Gene flow remains the same trend with the estimation under IM model but migration has increased in both directions compared with the IM model (Table 1). Consequently, Medium to high gene flow maintained in the speciation of the two pheasants in both categories with all models.

### Historical demography with PSMC

We sequenced and assembled the genome of *C. amherstiae* to serve as the reference. The genome assembly of *C. amherstiae* was 980Mb long (genomic coverage = 118.8×) and contained 21,996 scaffolds (N50 = 399Kb) (Table S8).

We conducted PSMC analyses to reconstruct the historical demography in both species of Phasianidae spanning the whole Pleistocene period, extending as far back as 5 million years (Fig. 3). The populations of two species of *Chrysolophus* were very similar in their estimated N_e_ history around 2 Mya, which is potentially consistent with the divergence time (2.23 Mya for allopatric population, 1.86 Mya for all sampled individuals) estimated by ABC analyses. At the start of the last ice age (110 thousand years ago, kya), the N_e_ of *C. pictus* increased slightly, and gradually decreased during the Last Interglacial (LIG), and ended with a sharp increase before the Last Glacial Maximum (LGM). From a peak about 30 kya, the N_e_ of *C. pictus* dropped by a factor of ten between 30 and 20 kya, throughout the LGM. Interestingly, the N_e_ of *C. amherstiae* showed a opposite trend to the N_e_ of *C. pictus*, decreasing from the last ice age to the beginning of LIG. We then saw a gradually increase in estimated N_e_ of *C. amherstiae* during LIG, and also a slightly growth in LGM.

**Fig. 3.**
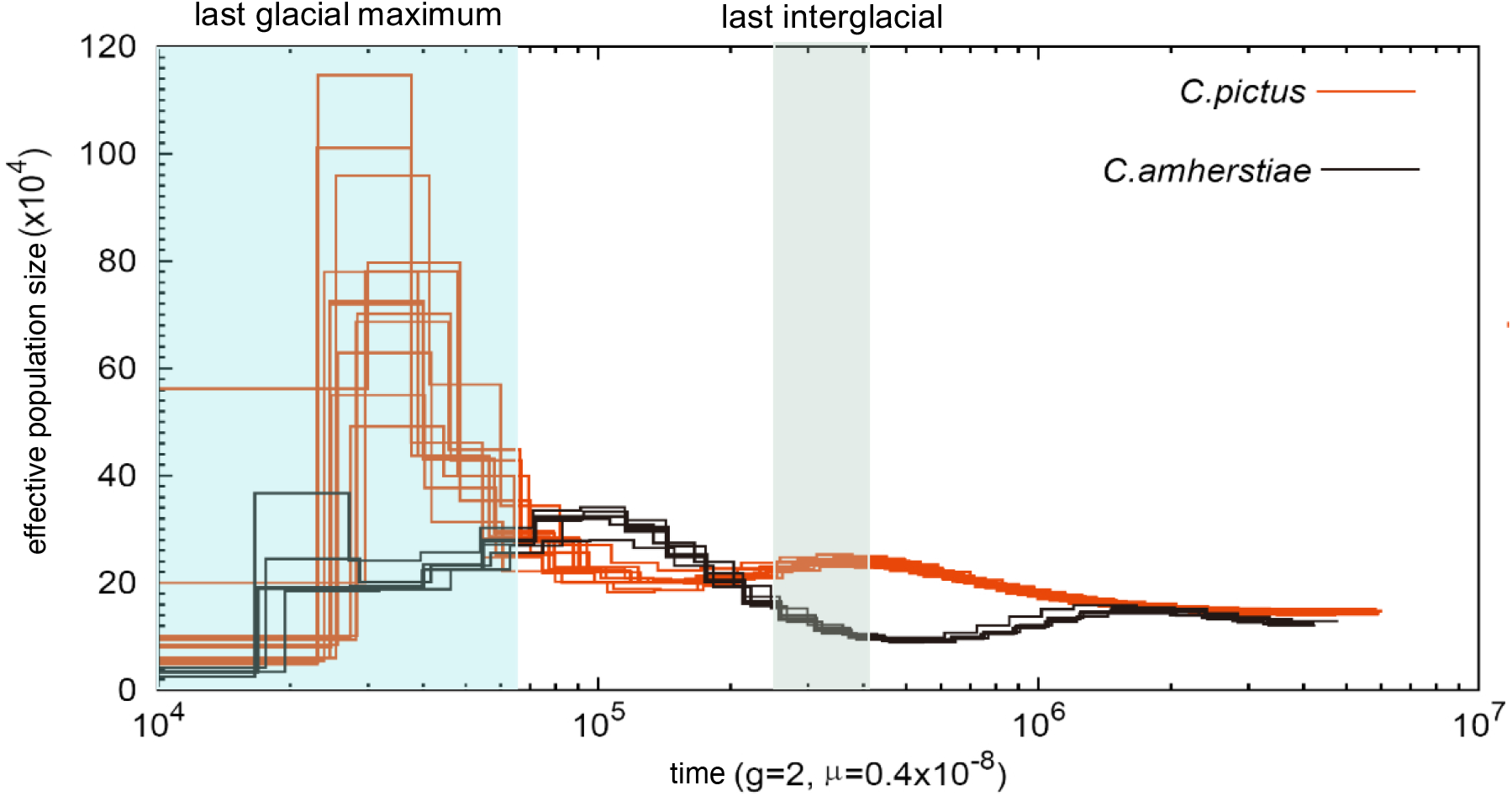
Inferred historical population sizes by Pairwise Sequential Markovian Coalescent analysis using 19 whole-genome sequenced individuals, including thirteen *C. pictus* and six *C. amherstiae* for re-sequencing data with an average of 35× depth. A *de novo* genome of *C. amherstiae* was used as the reference genome. The numbers of years per generation (g) for both species were set to be 2. The mutation rate per generation (μ) was set to 3.8 × 10^−9^ per site. (Orange: *C. pictus* Black: *C. amherstiae*)

## Discussion

Gene flow is conventionally considered as a homogenizing force that can moderate population divergence and impede speciation in the strict allopatric model of speciation. In this study, with multi-loci analysis by Approximate Bayesian Computation, we detected ancient and recurrent gene flow during the speciation process of the Golden pheasant and Lady Amherst’s pheasant, suggesting that the incomplete reproductive isolation has long existed between the two sexual dichromatic species. With genomic data-based PSMC analysis, we also show that the population expansion of Golden pheasant since Last Interglacial may played an essential impact in shaping the current suture zone and inducing hybridization in the wild.

### Ancient and recurrent gene flow

The evolution of reproductive isolation that underpins the Biological Species Concept allows us to reconsider the role of gene flow in speciation. Much of the studies implies that the types of reproductive isolation have been classified into two main categories: pre-zygotic isolation (e.g., assortative mating, sexual selection) and post-zygotic isolation (i.e., low hybrid fitness) (Coyne and Orr 2004; Nosil 2012). Gene flow during speciation may strengthen the barrier of reproductive isolation through reinforcement acts on pre-zygotic selection, as long as selection is not swamped by migration (Servedio and Kirkpatrick 1997; Kirkpatrick and Servedio 1999; Servedio 2000; Servedio 2004).

According to the results of ABC analyses for the allopatric populations of *Chrysolophus*, there has been long-term gene flow since the initial divergence of *C. pictus* and *C. amherstiae* two million years ago (Table 1). The ancient (or early) gene flow with speciation is not unusual in east Asia especially in the species complex distributed across the continental Taiwan Island and mainland China, e.g., the Hwamei (Li et al., 2010), bush warblers (Wei et al. 2019) and bamboo-partridges (Wang et al. 2021) or the closely related species diverged in the east edge of Qinghai-Tibet Plateau, e.g., the long-tailed tits (Wang et al. 2014), eared pheasants (Wang et al. 2017) and leaf-warblers (Zhang et al. 2019). The ecological divergent selection for local adaptation is often considered as the important factor that promotes ecological-driven speciation in the face of gene flow between these species without sexual dichromatic traits. Our results provided a new case that there was medium-level gene flow (Table 1) between the ancient populations of the two *Chrysolophus* species with significantly different ecological niches. However, the ecological segregation that limited the gene flow during the glacial time (see Lyu et al. 2015) may not be the major determinants for the separation of these two sexual dichromatic species.

The occasional hybrid records of Golden pheasants and Lady Amherst’s pheasants from the suture zone (see Deng 1974, He et al 1993, and Shi et al. 2018) and the less inviability of F_1_ hybrids or backcrosses in captivity (Arrieta et al. 2013) showed the probability that the current gene flow could be maintained due to the incompletely developed pre- and post-zygotic isolation. Our genetic data analysis revealed that when sympatric samples were included, the secondary contact model was better supported than when just allopatric samples were used. And the current high-level gene flow (Fig. 2) between the two pheasant species in the suture zone may have occurred after LIG (95% HPD: 0.13-1.206×10 years before) which is consistent with the estimation of expansion time based on the ecological niche model (see Lyu et al. 2015) and our PSMC estimation (Fig. 3). On the other side, we could not refuse the prediction of isolation with migration model with or without the sympatric samples (Table 1) allowing long-term gene flow since their divergence, which demonstrated the scenario of mixing-isolation-mixing process with recurrent gene flow should occur during their speciation. Since the two categories of factors that contributes to reproductive isolation, i.e., sexual dichromatic plumage which were usually attributed to the consequences of sexual selection that benefit to maintain the pre-zygotic isolation (Servedio and Noor 2003), and ecological segregation which may contribute more to the post-zygotic isolation (Funk et al. 2006), were both not completely developed between the Golden and Lady Amherst’s pheasant, with the fact that they were still able to maintain the independence of species level with divergent male plumage color and recurrent gene flow, we believe that the reinforcement selection on both sexual and ecological traits may play a significant role in the divergence and speciation of the two pheasant species. Further research into the fitness and attractiveness of the backcross or F_1_ hybrids in the suture zone would be helpful to undercover the mechanisms on the selection against the gene flow.

### Heterogeneous pattern of population demographic in LGM

It is widely perceived that climate change in the Pleistocene provided conditions for speciation (Hewitt 2000; Hewitt 2001). Gene flow might increase or decrease with climate change, depending on how climate change affects the habitat (Wasserman et al. 2010; Olsen et al. 2011). The cyclical climate changes in quaternary period (beginning 2.4 Mya), especially the LIG (between 130 kya and 70 kya) and LGM (between 26 kya and 18 kya), strongly affected the formation and distribution of species through “expansion-contraction” of their ranges (Hewitt 2000; Hewitt 2004; Provan and Bennett 2008). In East Asia, the impact of glacial period on the effective population size illustrated a different trend from that of high-latitude regions such as Europe and North America: For example, advancing ice sheets in the drove the expansion of polar biota and the contraction of temperate biota to refugia in glaciated regions such as Europe (Hewitt 2004; Provan and Bennett 2008), North America (Shafer et al. 2010) and Antarctica (Fraser et al. 2012). However, east Asia in the subtropical zone was not covered by ice sheets during the entire Pleistocene (Zhou et al. 2004), and the effective population size showed a contraction during the LIG and expansion during the LGM (Dong et al. 2017). According to the estimation of PSMC, the population dynamics of *C. pictus* and *C. amherstiae* were opposite, which can be inferred that LGM and LIG both had different effects on the two species, and the expansion of *C. pictus* during the recent secondary contact may provide opportunities for gene exchange (Fig. 3).

### Potential evolutionary processes contribute to reproductive isolation of the two sexual dichromatic pheasants

The evolutionary pattern of *Chrysolophus* belongs to the isolation with migration model, which means *C. pictus* and *C. amherstiae* have chances to encounter and hybridize during the evolutionary history. However, they haven’t fused into one species, instead maintained differentiation that significantly differ from phenotypes and ecological niches in the presence of long-term gene flow. Since the hybrids of the two pheasants were able to interbreed and produce fertile offspring, indicating that the post-zygotic isolation of *C. pictus* and *C. amherstiae* has not been fully established, and the barrier of reproductive isolation might be mainly improved by the promotion of pre-zygotic isolation (shown as sexual selection in our system). Each occurrence of gene flow is equivalent to a secondary contact, which increasingly strengthened the pre-zygotic isolation by reinforcement not only in the divergent male plumage color between the two pheasant species, but also may act on the cryptic divergence of female preference in male’s color which was beneficial to maintaining speciation with the recurrent gene flow.

## Conclusion

The results of the study support the scenario of speciation between Golden pheasant (*C. pictus*) and Lady Amherst’s pheasant (*C. amherstiae*) with cycles of mixing-isolation-mixing due to the dynamics of natural selection and sexual selection in late Pleistocene. This research provided a good research system as an evolutionary model to test reinforcement selection in speciation and to explore the genetics mechanisms for the origin and maintenance of reproductive isolation between sexual dichromatic species.

## Acknowledgments

We are grateful to Wanglang National Nature Reserve, Yanyun Zhang, Chao Zhao, Zhixin Zhou and Yiqiang Fu for their help in collecting samples. We thank Bowen Zhang, Yu Hao and Jindan Guo for assistance in statistics.

## Authors’ contributions

L.D. and J.Z. conceived the study; L.D. and W.L. collected the key samples; Z.L., J.Z. and M.G. performed the study and analyzed data; Z.L., J.Z. and L.D. drafted the manuscript with help from M.G. and W.L. All authors approved the final version of the manuscript.

## Funding

This study was supported by the National Natural Science Foundation of China (No.31471987).

## Availability of data and materials

The dataset used in the present study will be deposited to Genbank after the manuscript is accepted.

## Ethical note

The research protocol was approved by College of Life Sciences, Beijing Normal University: No. CLS-EAW-2013-007

## Consent for publication

Not applicable.

## Competing interests

The authors declare no conflicts of interests.

## Notes

### Competing Interest Statement

The authors have declared no competing interest.

