## Supplemental file for "Parapatric speciation with recurrent gene flow of two sexual dichromatic pheasants"

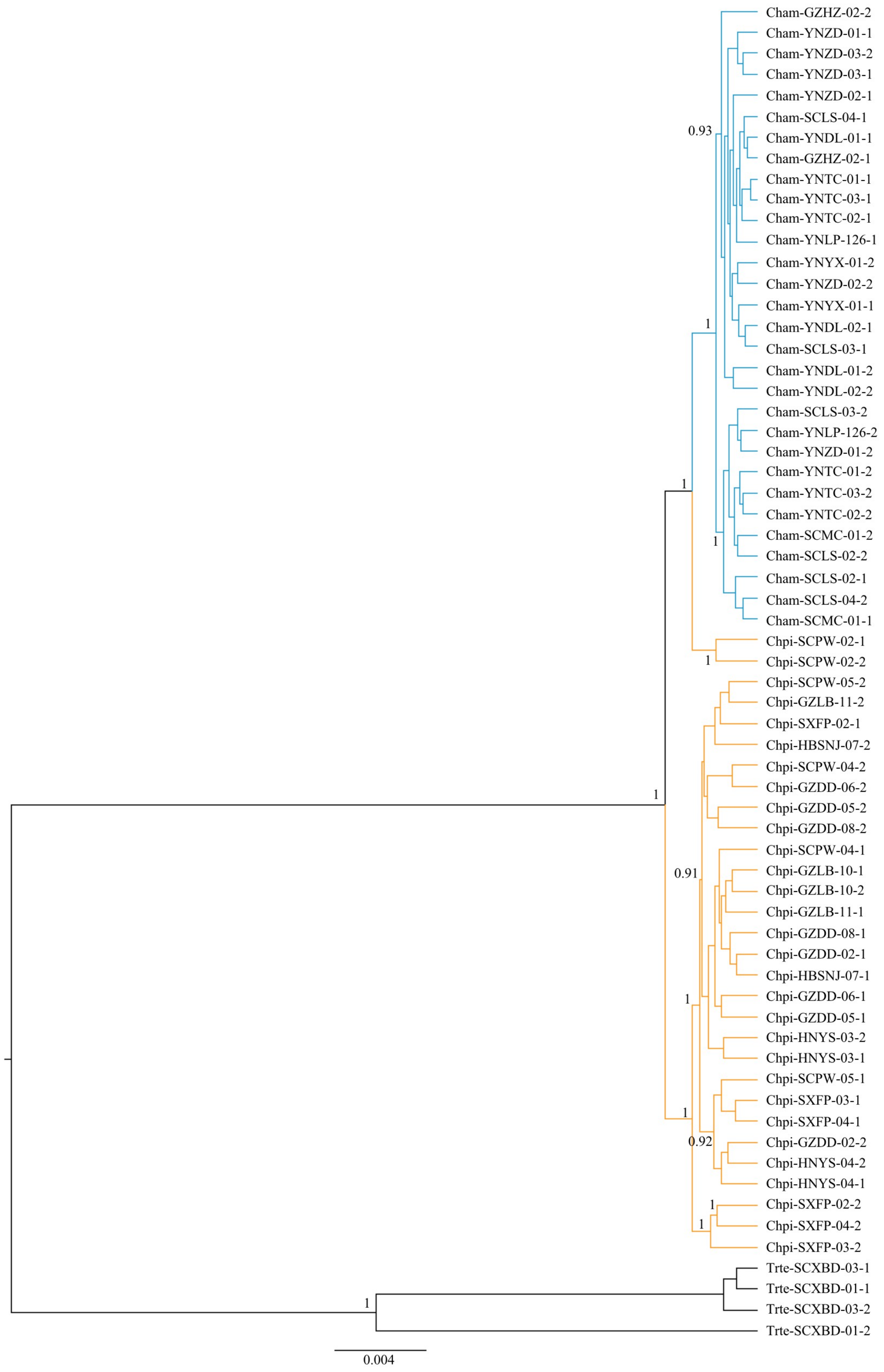


Figure S1. Bayesian tree reconstructed by autosome nuclear genes of Golden pheasants (orange) and Lady Amherst's pheasants (blue). The numbers on the branches are posterior probabilities above 0.9. Temminck's Tragopan *Tragopan temminckii* (black) was used as an outgroup. See Table S1 for abbreviations for each sampled individual.

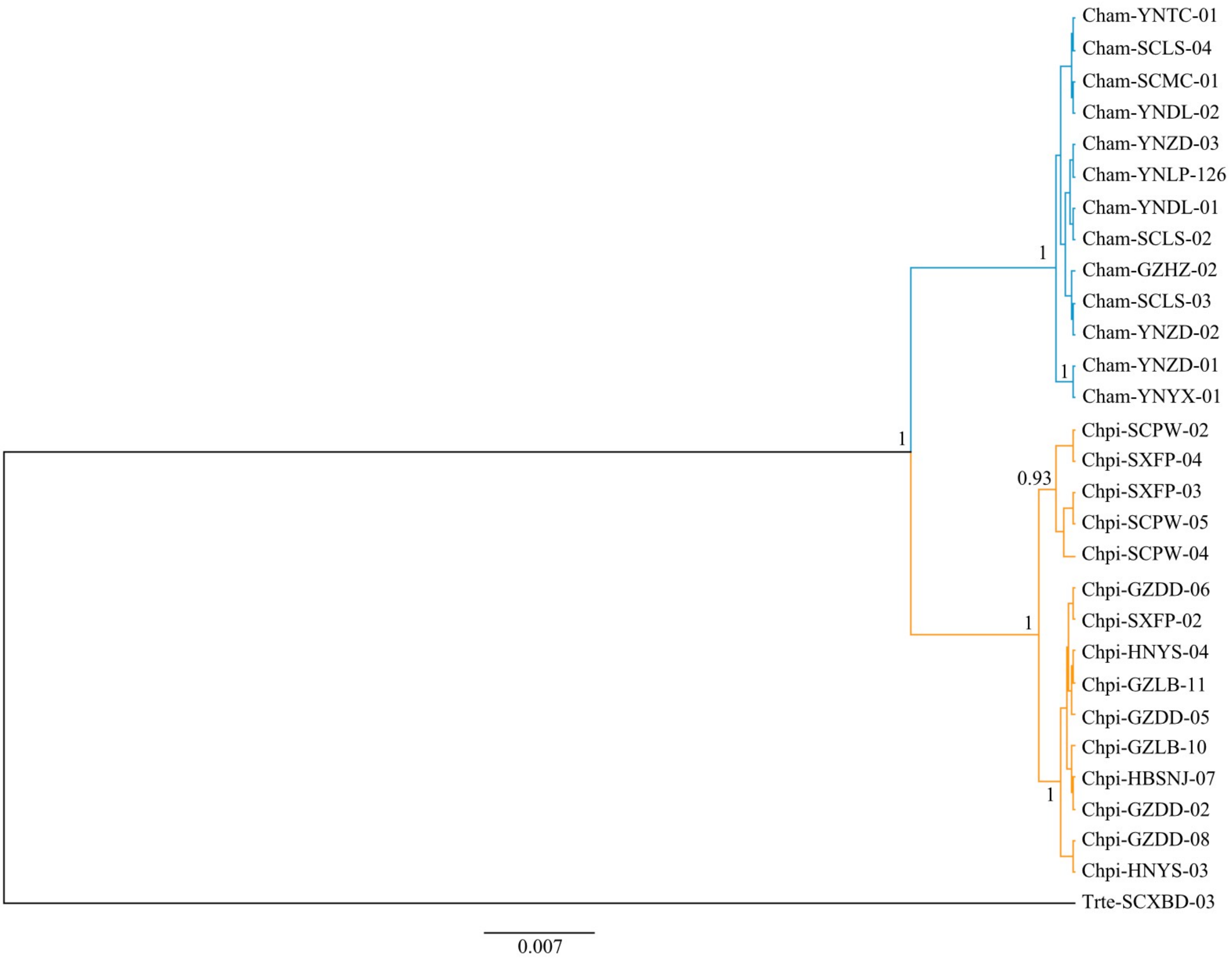


Figure S2. Bayesian tree reconstructed by mitochondria cyt *b* genes of Golden pheasants (orange) and Lady Amherst's pheasants (blue). The numbers on the branches are posterior probabilities above 0.9. Temminck's Tragopan *Tragopan temminckii* (black) was used as an outgroup. See Table S1 for abbreviations for each sampled individual.

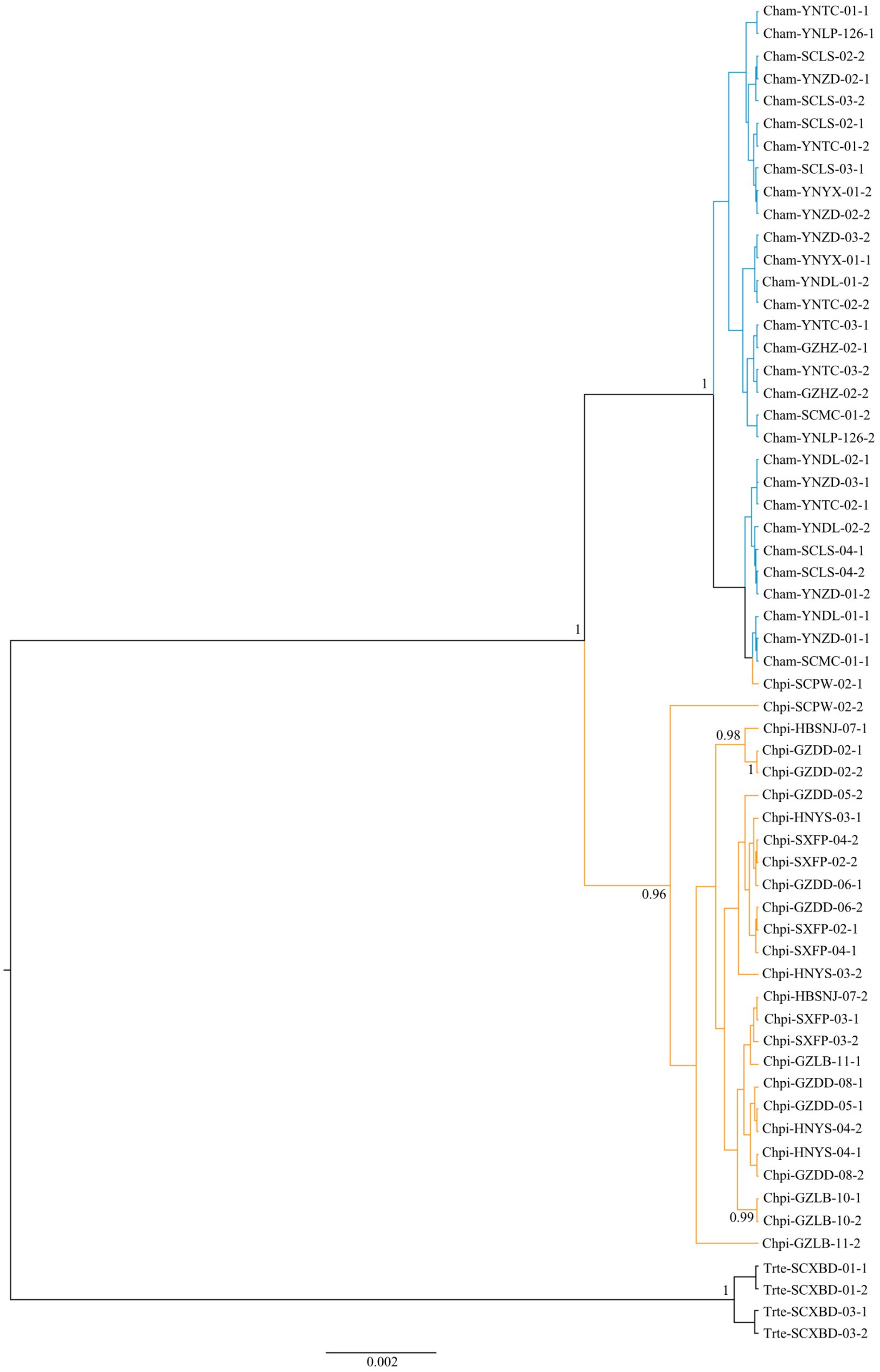


Figure S3. Bayesian tree reconstructed by three Z-link genes of Golden pheasants (orange) and Lady Amherst's pheasants (blue). The numbers on the branches are posterior probabilities above 0.9. Temminck's Tragopan *Tragopan temminckii* (black) was used as an outgroup. See Table S1 for abbreviations for each sampled individual.

Table S1 Sample information of *C. pictus* and *C. amherstiae*. The site number corresponds to the number in Figure 1. A total of 30 individuals were involved in the detection of hybrids using NewHybrids, and the hyphens represent the individuals have not used in the detection of hybrids.

| Species | Sample ID | Sampling site/Site number | For analysis | NewHybrids ID |
|---|---|---|---|---|
| *Chrysolophus pictus* | Chpi-GZDD-01 | Dadi, Guizhou / 5 | PSMC | - |
| | Chpi-GZDD-02 | Dadi, Guizhou / 5 | ABC | P1 |
| | Chpi-GZDD-05 | Dadi, Guizhou / 5 | ABC | P2 |
| | Chpi-GZDD-06 | Dadi, Guizhou / 5 | ABC, PSMC | P3 |
| | Chpi-GZDD-07 | Dadi, Guizhou / 5 | PSMC | - |
| | Chpi-GZDD-08 | Dadi, Guizhou / 5 | ABC, PSMC | P4 |
| | Chpi-GZLB-10 | Libo, Guizhou / 6 | ABC | P5 |
| | Chpi-GZLB-11 | Libo, Guizhou / 6 | ABC | P6 |
| | Chpi-HBSNJ-07 | Shennongjia, Hubei / 3 | ABC | P7 |
| | Chpi-HNYS-01 | Yongshun, Hunan / 4 | PSMC | - |
| | Chpi-HNYS-03 | Yongshun, Hunan / 4 | ABC | P8 |
| | Chpi-HNYS-04 | Yongshun, Hunan / 4 | ABC | P9 |
| | Chpi-SXFP-02 | Foping, Shanxi / 2 | ABC | P10 |
| | Chpi-SXFP-03 | Foping, Shanxi / 2 | ABC | P11 |
| | Chpi-SXFP-04 | Foping, Shanxi / 2 | ABC | P12 |
| | Chpi-SCPW-04 | Pingwu, Sichuan / 1 | ABC, PSMC | P13 |
| | Chpi-SCPW-05 | Pingwu, Sichuan / 1 | ABC, PSMC | P14 |
| | Chpi-SCPW-02 | Pingwu, Sichuan / 1 | ABC | P15 |
| | Chpi-SCPW-01 | Pingwu, Sichuan / 1 | PSMC | - |
| | Chpi-SCPW-03 | Pingwu, Sichuan / 1 | PSMC | - |
| | Chpi-CQ-01 | Wulong, Chongqing / 15 | PSMC | - |
| | Chpi-CQ-04 | Wulong, Chongqing / 15 | PSMC | - |
| *C. amherstiae* | Cham-GZHZ-02 | Hezhang, Guizhou / 9 | ABC, PSMC | A1 |
| | Cham-SCLS-01 | Leshan, Sichuan / 7 | PSMC | - |
| | Cham-SCLS-02 | Leshan, Sichuan / 7 | ABC | A2 |
| | Cham-SCLS-03 | Leshan, Sichuan / 7 | ABC | A3 |
| | Cham-SCLS-04 | Leshan, Sichuan / 7 | ABC | A4 |
| | Cham-SCMC-01 | Muchuan, Sichuan / 8 | ABC | A5 |
| | Cham-YNDL-01 | Dali, Yunnan / 11 | ABC, PSMC | A6 |
| | Cham-YNDL-02 | Dali, Yunnan / 11 | ABC, PSMC | A7 |
| | Cham-YNLP-126 | Lanping, Yunnan / 13 | ABC | A8 |
| | Cham-YNYX-01 | Yuxi, Yunnan / 10 | ABC, PSMC | A9 |
| | Cham-YNTC-01 | Tengchong, Yunnan / 12 | ABC | A10 |
| | Cham-YNTC-02 | Tengchong, Yunnan / 12 | ABC | A11 |
| | Cham-YNTC-03 | Tengchong, Yunnan / 12 | ABC | A12 |
| | Cham-YNZD-01 | Zhongdian, Yunnan / 14 | ABC, PSMC | A13 |
| | Cham-YNZD-02 | Zhongdian, Yunnan / 14 | ABC | A14 |
| | Cham-YNZD-03 | Zhongdian, Yunnan / 14 | ABC | A15 |

Table S2 The primer information of nuclear genes and cyt*b* gene.

| Number | Locus/LSH code | Primer | Primer sequence （5'→3'） |
|---|---|---|---|
| 1 | HERC1/10_63 | HERC1_F | ACTCATAAAGCAATCAAAGCTGA |
| | | HERC1_R | AATAAAGAAGTAGTGCCATTGCC |
| 2 | TMEM132B/15_3 | TMEM132B_F | GTAAACAACGATGCTACCAATCC |
| | | TMEM132B_R | AGGAGGGATCATGGACAGC |
| 3 | GPR15/1_70 | GPR15_F | GGGTCCAGAGGCTGATTGA |
| | | GPR15_R | CTCTGCGGATGTAGTTGTCAAAG |
| 4 | TTC28/15_24 | TTC28_F | ACAGACAAAAATATCCCCCAAGT |
| | | TTC28_R | CACCAGATGAAAGGTGACTATGA |
| 5 | KIAA1462/2_117 | KIAA1462_F | ACTGAGAGAGATAACACGCTTGG |
| | | KIAA1462_R | CAGGAGATGAGTGAAGACCAAAA |
| 6 | GPR125/4_80 | GPR125_F | AGCTTAGGAGCATTTTATGGACC |
| | | GPR125_R | TCCGAAACTGTACCTCTAATGGA |
| 7 | USP38/4_110 | USP38_F | CCTGAGATCCTTAGTGGAGACAA |
| | | USP38_R | GCATCCATGAGGTCTTTTTGTAG |
| 8 | DACT2/3_105 | DACT2_F | TCATCATAGTCAAGAGCAAGGCT |
| | | DACT2_R | GGTGGAGTGAAAATTAACAGCAG |
| 9 | DIDO1/20_47 | DIDO1_F | GTCAGAAAGGGAAACGTCCTG |
| | | DIDO1_R | GTTGTCTGAGGGCTTCTTCTTG |
| 10 | CEP350/8_3 | CEP350_F | TGCAAAGATGTTGATGACACTG |
| | | CEP350_R | TACAGCTGCTTCCCTTTCTAGC |
| 11 | CGNL1/10_16 | CGNL1_F | AATTTGCAGAAGTGGGAGTAGC |
| | | CGNL1_R | GGAGCCATATTTTGCTGGTTTTA |
| 12 | LOC100229969/3_24 | LOC100229969_F | AGAAAATGAACCCAGAAGCCAG |
| | | LOC100229969_R | GTGCACACTGTTGGCAGG |
| 13 | UBOX5/4_115 | UBOX5_F | CTGCGAGTGCTGGCTGAAG |
| | | UBOX5_R | GTTTACACCTCTACCTCATGCAA |
| 14 | SPECC1L/15_37 | SPECC1L_F | TTTGGCTATCAGTCTTTGAGTCC |
| | | SPECC1L_R | CAGCCTGTTCATCTCTATCTGGT |
| 15 | LOC100541159/11_26 | LOC100541159_F | CTCTGTCAAAGGACCTCAAAGC |
| | | LOC100541159_R | AGAATGACAAAGAATGGCAATGT |
| 16 | Z_62 | Z_62_F | CTAGCTCAGAAGCATCCAGGTC |
| | | Z_62_R | CAATTTCACTTTTGCTGTTTGGT |
| 17 | Z_86 | Z_86_F | CTGACTTGGGTACTGGAAGGTCT |
| | | Z_86_R | CCATTAAGCAAGAACCAGTGAA |
| 18 | 1_110 | 1_110_F | AATTGCATTCTTGATGGTGAAAT |
| | | 1_110_R | ACCACATCATATTTGTTGGAAGC |
| 19 | 1_117 | 1_117_F | GACCACAGGGACAGTATCCTACA |
| | | 1_117_R | TGAGATTTCTCCACTAAGATGGC |
| 20 | 1_191 | 1_191_F | ATCTGCAAGCTGTTCAGGC |
| | | 1_191_R | CCGGGCTGGGTTTTATTAC |

| | | | |
|---|---|---|---|
| 21 | 10_10 | 10_10_F | TGAGGCTAGAAGAAGTGAATGAAA |
| | | 10_10_R | ATTGTAGAAGGGCACACCTGAT |
| 22 | 10_19 | 10_19_F | AGTTCTCGTCAAACACCTGACA |
| | | 10_19_R | GCAAATACTTTCTCATTAGCCGA |
| 23 | 10_39 | 10_39_F1 | ATACTCATGTCATCCAAAAACCG |
| | | 10_39_R | CCCAACTTAATACCATCCAACAA |
| 24 | 11_15 | 11_15_F | CCAGTGTCTTCTACCTGATGGAG |
| | | 11_15_R | TTTGCACTTTTTCTATTTTCT |
| 25 | 14_42 | 14_42_F | GAACATCCTTACAGTTCTCAAAA |
| | | 14_42_R | TGGGGACTTCTTCCACTACAATA |
| 26 | 15_28 | 15_28_F | TCCATGATAACCTCAACCATTTC |
| | | 15_28_R | AACAACTACTACCCAGGGACACA |
| 27 | 21_3 | 21_3_F1 | CCAGTGAGATGGAGGCAGAG |
| | | 21_3_R | ATCCGCTCCCAGATGGC |
| 28 | 3_239 | 3_239_F | ATTTCCATTTATCCCATCCCATA |
| | | 3_239_R | CTTTTTGTGCCACTTCTGTAAGG |
| 29 | 3_68 | 3_68_F | GTTTGATTTTTGCTTCTCTGAGG |
| | | 3_68_R | GGCTGTGACCGTATTTATGCTAC |
| 30 | 4_108 | 4_108_F | ATCCACTCATCCCCTGGTC |
| | | 4_108_R | AAATGTTGCAGCACCCAAAC |
| 31 | 4_95 | 4_95_F | CTTTAGTCCACCACAGATTGGAC |
| | | 4_95_R | AGCTGCAAAGGGCATTCA |
| 32 | 6_29 | 6_29_F | AGTAGTTTCATTTTTCCAAGGGC |
| | | 6_29_R | CTTTGTTCCTGTTTACTGCCACT |
| 33 | CLTC | CLTC_F | CAGAATCCTGATCTAGCTTTACGAATGGC |
| | | CLTC_R | CATTTCTCCAGAAGTTGTTTGCGTCC |
| 34 | CLTCL1 | CLTCL1_F | CACCAATGTTCTGCAGAATCCTGA |
| | | CLTCL1_R | CCAGCTTATCTTCCTTNAGCCATTTCTC |
| 35 | FGB | FGB_F | CACGCCATATAGAGTATACTGTGACA |
| | | FGB_R | AACACTACCATCCTGGCGATTCTGAA |
| 36 | Serpinb14 | Serpinb14_F | GTTCGCTTTGATAAACTTCCAGG |
| | | Serpinb14_R | GGTGATTTGGTTGAGNATGTC |
| 37 | cyt *b* | AUKL | CCATCCAACATCTCAGCATGATGAAA |
| | | AUKH | GGAGTCTTCAGTCTCTGGTTTACAAGAC |

Table S3 Summary statistics of all nuclear genes. Length is the sequence length after alignment. N is the amount of the phased sequence that was used in the population genetic analysis. S is the number of polymorphic sites. $\Theta_\pi$ and $\Theta_w$ are two nucleotide diversity statistics. $F_{st}$ is the genetic statistic difference between *C. pictus* and *C. amhersitiae*. Hd is the haplotype diversity.

| Locus | Speices | Length (bp) | N | S | $\Theta_\pi$ | Hd | $\Theta_w$ | $F_{ST}$ |
|---|---|---|---|---|---|---|---|---|
| HERC1 | *C. pictus* | 628 | 24 | 6 | 0.00203 | 0.82971 | 0.00256 | 0.296 |
| HERC1 | *C. amhersitiae* | 628 | 28 | 6 | 0.00124 | 0.61905 | 0.00205 | |
| TMEM132B | *C. pictus* | 551 | 24 | 5 | 0.00088 | 0.612 | 0.00243 | 0.139 |
| TMEM132B | *C. amhersitiae* | 551 | 28 | 6 | 0.00174 | 0.86 | 0.0028 | |
| GPR15 | *C. pictus* | 577 | 24 | 1 | 0.00016 | 0.159 | 0.00046 | 0.874 |
| GPR15 | *C. amhersitiae* | 577 | 28 | 4 | 0.00042 | 0.384 | 0.00178 | |
| TTC28 | *C. pictus* | 567 | 24 | 10 | 0.00210 | 0.855 | 0.00472 | 0.585 |
| TTC28 | *C. amhersitiae* | 567 | 28 | 3 | 0.00040 | 0.373 | 0.00136 | |
| KIAA1462 | *C. pictus* | 721 | 24 | 5 | 0.00154 | 0.63 | 0.00188 | 0.196 |
| KIAA1462 | *C. amhersitiae* | 721 | 28 | 3 | 0.00100 | 0.585 | 0.00108 | |
| GPR125 | *C. pictus* | 584 | 24 | 3 | 0.00084 | 0.681 | 0.00138 | 0.624 |
| GPR125 | *C. amhersitiae* | 584 | 28 | 3 | 0.00064 | 0.563 | 0.00132 | |
| USP38 | *C. pictus* | 601 | 24 | 10 | 0.00191 | 0.746 | 0.00356 | 0.716 |
| USP38 | *C. amhersitiae* | 601 | 28 | 2 | 0.00014 | 0.14 | 0.00086 | |
| DACT2 | *C. pictus* | 823 | 24 | 12 | 0.00254 | 0.826 | 0.00392 | 0.489 |
| DACT2 | *C. amhersitiae* | 823 | 28 | 7 | 0.00156 | 0.757 | 0.00219 | |
| DIDO1 | *C. pictus* | 734 | 24 | 11 | 0.00267 | 0.862 | 0.00401 | 0.673 |
| DIDO1 | *C. amhersitiae* | 734 | 28 | 2 | 0.00032 | 0.315 | 0.0007 | |
| CEP350 | *C. pictus* | 587 | 24 | 8 | 0.00205 | 0.866 | 0.00381 | 0.566 |
| CEP350 | *C. amhersitiae* | 587 | 28 | 10 | 0.00314 | 0.868 | 0.00456 | |
| CGNL1 | *C. pictus* | 620 | 24 | 4 | 0.00048 | 0.301 | 0.00175 | 0.819 |
| CGNL1 | *C. amhersitiae* | 620 | 28 | 8 | 0.00129 | 0.701 | 0.00336 | |
| LOC100229969 | *C. pictus* | 645 | 24 | 11 | 0.00223 | 0.772 | 0.00457 | 0.737 |
| LOC100229969 | *C. amhersitiae* | 645 | 30 | 4 | 0.00020 | 0.526 | 0.00157 | |
| UBOX5 | *C. pictus* | 528 | 24 | 12 | 0.00319 | 0.848 | 0.00609 | 0.564 |
| UBOX5 | *C. amhersitiae* | 528 | 28 | 8 | 0.00154 | 0.746 | 0.00391 | |
| SPECC1L | *C. pictus* | 743 | 22 | 14 | 0.00049 | 0.939 | 0.0052 | 0.381 |
| SPECC1L | *C. amhersitiae* | 732 | 28 | 12 | 0.00134 | 0.521 | 0.00421 | |
| LOC100541159 | *C. pictus* | 559 | 24 | 11 | 0.00383 | 0.909 | 0.00527 | 0.291 |
| LOC100541159 | *C. amhersitiae* | 559 | 28 | 8 | 0.00297 | 0.839 | 0.00368 | |
| Z-62 | *C. pictus* | 667 | 22 | 6 | 0.00096 | 0.593 | 0.00247 | 0.877 |
| Z-62 | *C. amhersitiae* | 667 | 26 | 1 | 0.00049 | 0.492 | 0.00039 | |
| Z-86 | *C. pictus* | 627 | 24 | 5 | 0.00078 | 0.493 | 0.00214 | 0.829 |
| Z-86 | *C. amhersitiae* | 627 | 24 | 1 | 0.00043 | 0.431 | 0.00044 | |
| 1_110 | *C. pictus* | 1017 | 24 | 6 | 0.00195 | 0.757 | 0.00158 | 0.334 |
| 1_110 | *C. amhersitiae* | 1017 | 28 | 3 | 0.00067 | 0.585 | 0.00076 | |
| 1_117 | *C. pictus* | 799 | 22 | 6 | 0.00199 | 0.71 | 0.00206 | 0.541 |
| 1_117 | *C. amhersitiae* | 799 | 28 | 5 | 0.00110 | 0.717 | 0.00161 | |
| 1_191 | *C. pictus* | 538 | 24 | 5 | 0.00128 | 0.739 | 0.00249 | 0.284 |
| 1_191 | *C. amhersitiae* | 538 | 28 | 2 | 0.00065 | 0.579 | 0.00096 | |
| 10_10 | *C. pictus* | 681 | 24 | 9 | 0.00191 | 0.797 | 0.00354 | 0.476 |

| | | | | | | | | |
|---|---|---|---|---|---|---|---|---|
| 10_10 | *C. amhersitiae* | 681 | 28 | 4 | 0.00139 | 0.69 | 0.00151 | |
| 10_19 | *C. pictus* | 615 | 24 | 6 | 0.00164 | 0.783 | 0.00261 | 0.692 |
| 10_19 | *C. amhersitiae* | 615 | 28 | 2 | 0.00014 | 0.14 | 0.00084 | |
| 10_39 | *C. pictus* | 619 | 24 | 8 | 0.00213 | 0.851 | 0.00346 | 0.178 |
| 10_39 | *C. amhersitiae* | 619 | 28 | 5 | 0.00112 | 0.545 | 0.00208 | |
| 11_15 | *C. pictus* | 640 | 24 | 2 | 0.00060 | 0.554 | 0.00084 | 0.62 |
| 11_15 | *C. amhersitiae* | 640 | 28 | 5 | 0.00097 | 0.519 | 0.00201 | |
| 14_42 | *C. pictus* | 844 | 24 | 13 | 0.00463 | 0.891 | 0.00411 | 0.215 |
| 14_42 | *C. amhersitiae* | 844 | 28 | 9 | 0.00180 | 0.796 | 0.00278 | |
| 15_28 | *C. pictus* | 584 | 24 | 15 | 0.00284 | 0.906 | 0.00691 | 0.213 |
| 15_28 | *C. amhersitiae* | 584 | 28 | 4 | 0.00092 | 0.675 | 0.00177 | |
| 21_3 | *C. pictus* | 564 | 24 | 10 | 0.00321 | 0.888 | 0.00475 | 0.428 |
| 21_3 | *C. amhersitiae* | 564 | 28 | 10 | 0.00171 | 0.709 | 0.00456 | |
| 3_239 | *C. pictus* | 679 | 24 | 10 | 0.00384 | 0.862 | 0.00403 | 0.375 |
| 3_239 | *C. amhersitiae* | 679 | 24 | 7 | 0.00130 | 0.761 | 0.0032 | |
| 3_68 | *C. pictus* | 647 | 24 | 10 | 0.00216 | 0.833 | 0.00414 | 0.594 |
| 3_68 | *C. amhersitiae* | 647 | 28 | 3 | 0.00089 | 0.64 | 0.00119 | |
| 4_108 | *C. pictus* | 564 | 24 | 5 | 0.00178 | 0.801 | 0.00239 | 0.182 |
| 4_108 | *C. amhersitiae* | 564 | 26 | 4 | 0.00125 | 0.668 | 0.00187 | |
| 4_95 | *C. pictus* | 607 | 24 | 5 | 0.00178 | 0.757 | 0.00221 | 0.54 |
| 4_95 | *C. amhersitiae* | 607 | 28 | 3 | 0.00107 | 0.664 | 0.00127 | |
| 6_29 | *C. pictus* | 737 | 24 | 3 | 0.00086 | 0.685 | 0.00109 | 0.772 |
| 6_29 | *C. amhersitiae* | 737 | 28 | 5 | 0.00096 | 0.598 | 0.00174 | |
| CLTC | *C. pictus* | 852 | 24 | 11 | 0.00349 | 0.873 | 0.00346 | 0.312 |
| CLTC | *C. amhersitiae* | 852 | 28 | 12 | 0.00494 | 0.862 | 0.00362 | |
| CLTCL1 | *C. pictus* | 486 | 24 | 1 | 0.00022 | 0.29 | 0.00108 | 0.913 |
| CLTCL1 | *C. amhersitiae* | 486 | 26 | 2 | 0.00022 | 0.218 | 0.00108 | |
| FGB | *C. pictus* | 351 | 24 | 6 | 0.00248 | 0.703 | 0.00458 | 0.456 |
| FGB | *C. amhersitiae* | 351 | 24 | 5 | 0.00152 | 0.656 | 0.00384 | |
| Serpinb14 | *C. pictus* | 557 | 24 | 6 | 0.00221 | 0.728 | 0.00301 | 0.413 |
| Serpinb14 | *C. amhersitiae* | 557 | 26 | 6 | 0.00098 | 0.569 | 0.00294 | |

Table S4 Summary statistics of neutrality test of 37 genes. Both $\lambda^2$(HKA) and Tajima's D are considered significant at P<0.01 (shown as "**").

| Number | Locus/LSH code | $\lambda^2$(HKA) | Tajima's D |
|---|---|---|---|
| 1 | HERC1/10_63 | 0.339 | 0.02386 |
| 2 | TMEM132B/15_3 | 0.042 | -0.91812 |
| 3 | GPR15/1_70 | 0.262 | -0.40341 |
| 4 | TTC28/15_24 | 0.005 | -1.65764 |
| 5 | KIAA1462/2_117 | 0.019 | -0.53983 |
| 6 | GPR125/4_80 | 2.636 | -0.32844 |
| 7 | USP38/4_110 | 1.495 | -0.47015 |
| 8 | DACT2/3_105 | 0.162 | -1.04148 |
| 9 | DIDO1/20_47 | 0.673 | 0.05655 |
| 10 | CEP350/8_3 | 0.112 | 0.23184 |
| 11 | CGNL1/10_16 | 0.028 | -0.20575 |
| 12 | LOC100229969/3_24 | 0.096 | -1.06589 |
| 13 | UBOX5/4_115 | 1.166 | -0.08835 |
| 14 | SPECC1L/15_37 | 0.239 | -0.79036 |
| 15 | LOC100541159/11_26 | 0.347 | 0.67994 |
| 16 | Z_62 | 0.303 | 0.99660 |
| 17 | Z_86 | 2.568 | 0.43554 |
| 18 | 1_110 | 0.743 | -0.59105 |
| 19 | 1_117 | 0.002 | -0.28749 |
| 20 | 1_191 | 0.077 | -0.77047 |
| 21 | 10_10 | 1.082 | -0.64961 |
| 22 | 10_19 | 2.949 | 0.09397 |
| 23 | 10_39 | 0.577 | -0.91940 |
| 24 | 11_15 | 2.037 | -0.19945 |
| 25 | 14_42 | 0.135 | -1.08822 |
| 26 | 15_28 | 0.056 | -1.59917 |
| 27 | 21_3 | 0.038 | 0.03041 |
| 28 | 3_239 | 0.126 | -0.46038 |
| 29 | 3_68 | 0.775 | -0.06200 |
| 30 | 4_108 | 0.725 | -0.07365 |
| 31 | 4_95 | 1.085 | 0.83198 |
| 32 | 6_29 | 0.177 | 0.63547 |
| 33 | CLTC | 0.325 | 0.78896 |
| 34 | CLTCL1 | 0.138 | 0.07042 |
| 35 | FGB | 0.005 | 0.32964 |
| 36 | Serpinb14 | 0.057 | -0.76561 |
| 37 | cyt *b* | 1.040 | 2.64308** |

Table S5 The prior range of each parameter in msABC analysis. $N_{cp}$: effective population size of *C. pictus*; $N_{ca}$: effective population size of *C. amherstiae*; $N_A$: effective population size of the common ancestor of *Chrysolophus*; $T_{div}$: population divergence time; $T_1$: Termination time of early gene flow; $T_2$: termination time of gene flow after secondary contact; $M_1$: the number of migrants from *C. amherstiae* to *C. pictus* in each generation. $M_2$: the number of migrants from *C. pictus* to *C. amherstiae* in each generation.

| Speciation model | Effective population size（$10^5$） | | | Time（$10^5$ years） | | | Migrants | |
|---|---|---|---|---|---|---|---|---|
| | $N_{cp}$ | $N_{ca}$ | $N_A$ | $T_1$ | $T_2$ | $T_{div}$ | $M_1$ | $M_2$ |
| Isolation model | 0-5 | 0-5 | 0-10 | —— | —— | 0-5 | 0 | 0 |
| Isolation with migration model | 0-5 | 0-5 | 0-10 | —— | —— | 0-5 | 0-30 | 0-30 |
| Early gene flow model | 0-5 | 0-5 | 0-10 | 0.1-5 | —— | 0-5 | 0-30 | 0-30 |
| Secondary contact model | 0-5 | 0-5 | 0-10 | —— | 0-0.2 | 0-5 | 0-30 | 0-30 |

Table S6 Command line for each model in msABC. The meaning of each parameter can be found in the instruction of the msABC.

| Demorgraphic model | msABC command |
|---|---|
| Isolation | ./msABC 52 1000000 --dur-mode -I 2 24 28 -n 1 -U 0 5 -n 2 -U 0 5 -ej -U 0 5 2 1 -eN 0 -U 0 10 --frag-begin --finp locfile.txt --N 100000 --frag-end --verbose > out.txt |
| Isolation with migration | ./msABC 52 1000000 --dur-mode -I 2 24 28 -m 1 2 -U 0 30 -m 2 1 -U 0 30 -n 1 -U 0 5 -n 2 -U 0 5 -ej -U 0 5 2 1 -eN 0 -U 0 10 --frag-begin --finp locfile.txt --N 100000 --frag-end --verbose > out.txt |
| Early gene flow model | ./msABC 52 1000000 --dur-mode -I 2 24 28 -m 1 2 0 -m 2 1 0 -n 1 -U 0 5 -n 2 -U 0 5 -em -U 0.1 5 1 2 -U 0 30 -em 0 2 1 -U 0 30 -ej -U 0 5 2 1 -eN 0 -U 0 10 --frag-begin --finp locfile.txt --N 100000 --frag-end --verbose >out.txt |
| Secondary contact model | ./msABC 52 1000000 --dur-mode -I 2 24 28 -m 1 2 -U 0 30 -m 2 1 -U 0 30 -n 1 -U 0 5 -n 2 -U 0 5 -em -U 0 0.2 1 2 0 -em 0 2 1 0 -ej -U 0 5 2 1 -eN 0 -U 0 10 --frag-begin --finp locfile.txt --N 100000 --frag-end --verbose > out.txt |

Table S7. The sequencing data of the *de novo C. amherstiae* genome.

| | Length of DNA insertion fragment | Data size（G） | Length of paired-end sequencing（bp） | Depth（X） |
|---|---|---|---|---|
| Sequencing data | 450bp | 62.4 | 250 | 63.7 |
| | 5K | 54 | 150 | 55.1 |
| Total | — | 116.4 | — | 118.8 |

Table S8 Assembly results of the *C. amherstiae* genome.

| | Length | | Count | |
|---|---|---|---|---|
| | Contig（bp） | Scaffold（bp） | Contig（bp） | Scaffold（bp） |
| Total | 997,601,024 | 1,038,879,668 | 192,791 | 21,996 |
| maximum | 967,879 | 6,872,762 | — | — |
| Count>=2000 | — | — | 131,539 | 12,455 |
| N50 | 13,040 | 399,346 | 14,743 | 638 |
| N60 | 5,829 | 286,160 | 25,907 | 943 |
| N70 | 3,458 | 203,102 | 50,264 | 1,370 |
| N80 | 2,718 | 123,646 | 82,933 | 2,021 |
| N90 | 2,089 | 50,805 | 124,721 | 3,304 |
